# Maternal obesity increases hypothalamic miR-505-5p expression in mouse offspring leading to altered fatty acid sensing and increased intake of high-fat food

**DOI:** 10.1101/2022.06.01.494310

**Authors:** Laura Dearden, Isadora C. Furigo, Lucas C. Pantaleão, L W. P. Wong, Denise S. Fernandez-Twinn, Juliana de Almeida-Faria, Katherine A. Kentistou, Maria V. Carreira, Guillaume Bidault, Antonio Vidal-Puig, Ken K. Ong, John R. B. Perry, Jose Donato, Susan E. Ozanne

## Abstract

*In utero* exposure to maternal obesity programs increased obesity risk. Animal models show that programmed offspring obesity is preceded by hyperphagia, but the mechanisms that mediate these changes are unknown. Using a mouse model of maternal obesity, we observed increased intake of a high-fat diet in offspring of obese mothers that precedes the development of obesity. Through small RNA sequencing, we identified programmed overexpression of hypothalamic miR-505-5p that is established in the fetus, lasts to adulthood and is maintained in hypothalamic neural progenitor cells cultured *in vitro*. Metabolic hormones and long-chain fatty acids associated with obesity increase miR-505-5p expression in hypothalamic neurons *in vitro*. We demonstrate that targets of miR-505-5p are enriched in fatty acid metabolism pathways and over-expression of miR-505-5p decreased neuronal fatty acid metabolism *in vitro*. miR-505-5p targets are associated with increased BMI in human genetic studies. Intra-cerebroventricular injection of miR-505-5p in wild-type mice increased high-fat diet intake, mimicking the phenotype observed in offspring exposed to maternal obesity. Conversely, maternal exercise intervention in an obese mouse pregnancy rescued the programmed increase of hypothalamic miR-505-5p in offspring of obese dams and reduced high-fat diet intake to control offspring levels. This study identifies a novel mechanism by which maternal obesity programs obesity in offspring via increased intake of high-fat foods.

## 1. Introduction

The rapid global rise in obesity represents a significant health challenge. A genetic influence on body weight is undisputed, but the recent rapid increase in obesity prevalence suggests a robust interplay between genetic factors and the environment (1). It is now well established that an individual’s long-term health can be influenced by *in utero* and early post-natal environmental conditions, giving rise to the concept of “developmental programming” (2). Numerous studies in humans and animal models have highlighted the association between maternal obesity and several long-term adverse health outcomes in the offspring, including the risk of obesity and type 2 diabetes (3). Growing evidence from animal models shows that exposure to maternal obesity can disrupt the hypothalamic circuits responsible for maintaining energy homeostasis, which may contribute to increased offspring susceptibility to obesity later in life (4).

The hypothalamus is the crucial region of the brain responsible for maintaining energy homeostasis in the body and contains specialised neurons that sense circulating nutrients and hormones. Exposure to over-nutrition during critical windows of development has been shown to program increased food intake in animal models (4), including increased preference for high-fat foods in rats and non-human primates (5–7). Studies in mouse models report that maternal obesity alters hypothalamic neuronal projections (8–10), but aside from this little is known about the molecular mechanisms by which maternal overnutrition programs permanent changes in feeding behaviour in offspring.

The stable nature of metabolic phenotypes in offspring exposed to maternal obesity during development suggests lasting changes to gene profiles across the lifespan. There is accumulating evidence that miRNAs, which are small non-coding RNAs that regulate gene function through degradation of mRNAs and/or inhibition of protein translation (11), may reflect an important mechanism by which the maternal environment can alter long-term phenotypes in the offspring (3, 12). We have reported previously that programmed cell-autonomous up-regulation of miR-126 in adipose tissue of offspring of obese mouse dams is responsible for the downregulation of the genes *Irs1* and *Lnpk*, contributing to insulin resistance and endoplasmic reticulum stress, respectively, in these animals (13, 14). Few studies to date have explored the role of miRNAs in regulating hypothalamic energy-sensing (15–19), or examined the potential contribution of miRNAs to programming of long-term hypothalamic function by the perinatal environment (20).

A growing body of research suggests that neurons within the hypothalamus can sense and respond to changes in fatty acid (FA) levels in the circulation (21–24), as reported previously for glucose-sensing neurons (25). Circulating FAs, particularly long-chain fatty acids, may act as cellular messengers, which inform sub-populations of neurons in the hypothalamus about whole body nutrient status, allowing the regulation of food intake, hepatic glucose production and insulin secretion in order to maintain energy homeostasis (22, 26, 27). Studies in rodents and humans have shown that oleic and palmitic acid reduce the expression of neuropeptide Y (NPY) and increase expression of pro-opiomelanocortin (POMC)(28–30), and are associated with a reduction in food intake. Furthermore, both oleic (31) and octanoic acid (32) can reduce food intake and increase energy expenditure via regulation of POMC neuron excitability, confirming that several FAs can act centrally to regulate food intake. Notably, some studies have suggested that neuronal responses to FAs are blunted in diet-induced obese animals (33, 34), and this blunting of neuronal FA sensing may contribute to unbalanced energy homeostasis. Whether the nutritional environment in early life impacts long-term neuronal FA sensing-as has been reported for glucose sensing (35, 36)-remains unclear. However, one study has suggested that maternal HFD consumption during pregnancy alters the sensitivity of offspring ventromedial hypothalamic neurons to oleic acid (35).

Therefore, this study aimed to explore the contribution of hypothalamic miRNAs in mediating the effects of maternal obesity on offspring obesity risk. We profiled miRNA expression in the offspring hypothalamus to identify miRNAs that could mediate increased high-fat diet intake and adopted a combination of *in vitro* and *in vivo* approaches to demonstrate the causal effects of altered levels of a specific miRNA on hypothalamic neuronal function and feeding behaviour.

## 2. Methods

### 2.1. Mouse model

We utilised a mouse model of maternal diet-induced obesity published previously by our group (37). Briefly, female C57BL/6J mice were randomly assigned prior to mating to either a control chow diet (RM1; Special Dietary Services, Witham UK) or a highly palatable obesogenic high-fat diet (45% total calories from fat, Special Dietary Services, Witham UK) supplemented with condensed milk (Nestle, UK) fortified with mineral mix AIN93G (Special Dietary Services, Witham UK). After approximately three weeks on the respective diets, females were mated with chow-fed males for their first pregnancy. Dams remained on their respective diets throughout pregnancy and lactation and the first litter was culled after weaning. After first pregnancy, proven breeders from the obese group when they had achieved a body composition of around 30–35% fat and body weight exceeding 35g were mated for second pregnancy along with age matched lean dams. A subset of dams was culled on embryonic day 13 (E13) to collect fetal hypothalamus and remaining pregnancies were allowed to deliver naturally. On post-natal day 2 litters were reduced to 6 pups to ensure standardized milk supply (38). One male and female mouse per litter was included in each separate experiment, therefore the statistical number relates to the dam. For diet-induced obesity experiments in adult male mice in Figure 2k, mice were fed *ad-libitum* chow diet (DS-105, Safe diets) or HFD (60% total calories from fat, D12492, Research diets) for 21 weeks.

### 2.2. Food intake and body composition analyses in offspring

At weaning, male and female mice from control or obese mothers were randomly assigned to either a control chow (as above) or obesogenic diet (as above), creating 4 experimental offspring groups: offspring born to control mothers weaned onto a chow diet (Off-C Chow), offspring born to control mothers weaned onto a HFD (Off-C HFD), offspring born to obese mothers weaned onto a chow diet (Off-Ob Chow), offspring born to obese mothers weaned onto a HFD (Off-Ob HFD) (illustrated in Figure 1A). *Ad libitum* food intake in the home cage was measured twice a week from the age of 3 weeks to 8 weeks. Animals were weighed weekly, and body composition determined in fed animals using Time-Domain Nuclear Magnetic Resonance imaging (TD-NMR; Minispec Plus, Bruker, Billerica, MA, USA).

**Fig 1.**
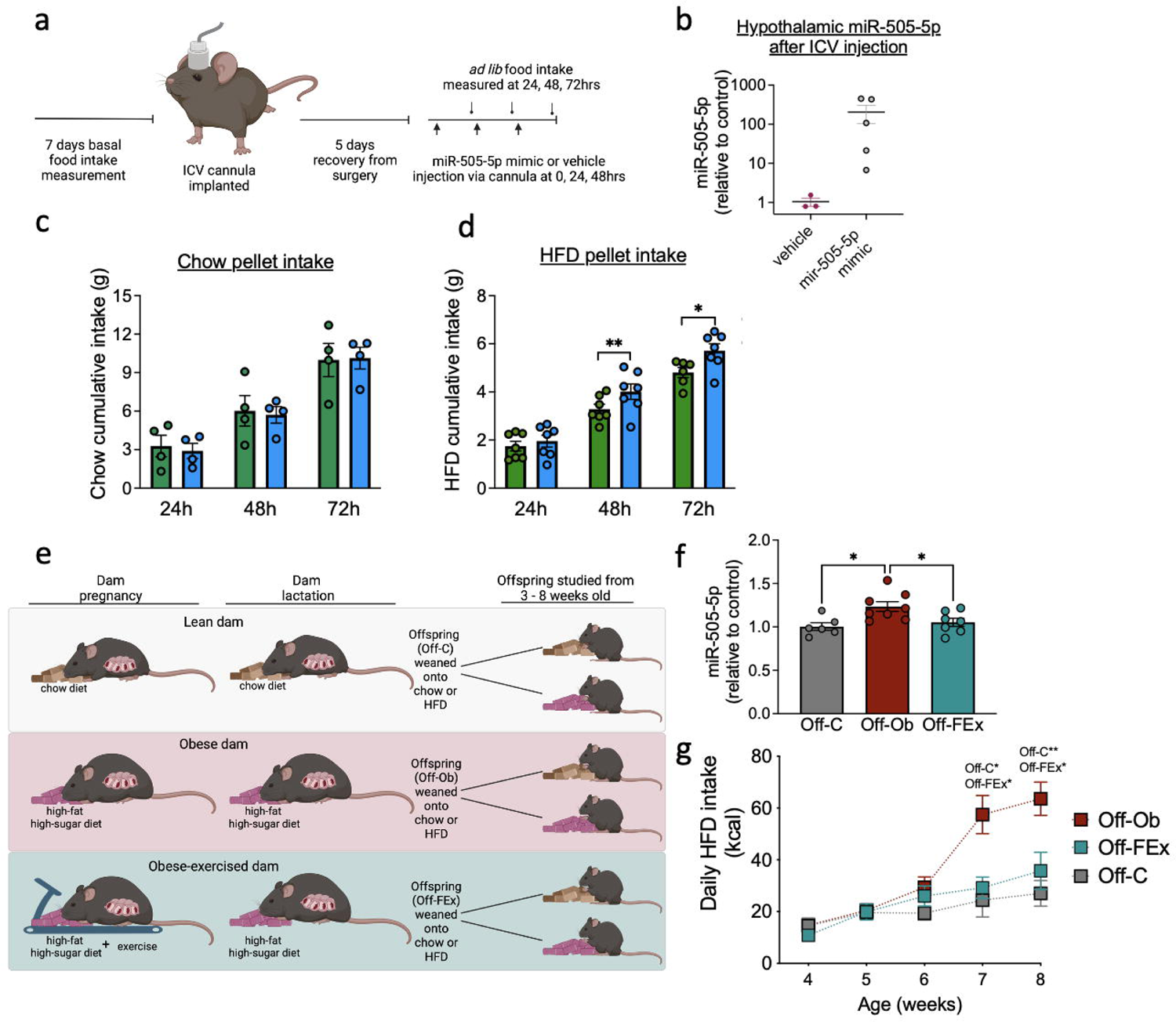
Offspring of obese mothers eat more of a high-fat pellet and develop increased adiposity compared to offspring of lean mothers fed the same diet. a) Diagram depicting mouse model of maternal obesity. At weaning offspring are weaned onto either a chow or HFD, and food intake, body weight and adiposity are measured weekly. b) Normal chow daily kilocalorie intake of male offspring born to control (Off-C; n=21) and obese mothers (Off-Ob; n=15). c) High fat diet (HFD) pellet daily kilocalorie intake of male offspring born to control (Off-C; n=22) and obese mother (Off-Ob; n=19). d) Sucrose preference of male offspring during a two-bottle choice test (Off-C n=6, Off-Ob n=4). e) Intake (g) of sucrose solution during two bottle choice test (Off-C n=6, Off-Ob n=4). f) Body weight and (g) fat mass from 4 to 8 weeks of age in offspring weaned onto a chow diet (Off-C n=12, Off-Ob n=21). h) Body weight and (i) fat mass from 4 to 8 weeks of age in offspring weaned onto a high-fat diet (Off-C n=12 Off-Ob n=13). **P<0.01, ***P<0.001 Statistical significance was determined using mixed effects analysis with Sidak’s post-hoc test. The underlying data are provided as a Source Data file

For two-bottle choice tests for sucrose mice were single housed at six weeks of age. At eight weeks of age, baseline measurements of body weight, chow intake and water intake were measured daily for three days. There were no differences in baseline measurements. A sucrose preference test was conducted across a range of sucrose solutions −0.5%, 1%, 2%, 4%, 8%, 16% and 32% [w/v] in water. Mice were habituated to the set-up for three days, with one bottle containing 0.5% sucrose ([w/v] in drinking water) and the other water, during which body weight, chow intake, and the weight of the water and sucrose bottles were measured daily at 1300. Bottles were weighed before they were placed in the top of the cages. After 24 hours, the bottles were carefully removed and weighed again. The side order of the bottles was switched daily to avoid side preference.

### 2.1. Exercise intervention in dams

From one week prior to mating for the second pregnancy, a group of obese dams underwent an exercise intervention protocol that we have published previously (39). Briefly, dams were trained to run 20lllminutes daily on a treadmill, starting at 5lllm/min on the first day and with daily incremental increases in speed until they were running at a final speed of 12.5lllm/min on day 5. Dams were then rested for 2 days during the weekend when they were time-mated and day 1 of pregnancy was defined by the appearance of a copulatory plug. Exercise was then maintained at the final running speed and following a routine of 5 days of daily exercise (weekdays) followed by 2 days of rest (weekends), until day 17. The duration of running at the top speed of 12.5lllm/min was reduced in the final 3 days to adapt to the physical constraints of pregnancy. At weaning, half of the offspring were weaned onto an obesogenic HFD (as above-to assess the impact of the intervention on feeding behaviour) and half were weaned onto a chow diet (to examine hypothalamic gene expression independent of the effects of an HFD).

### 2.2. Small RNA sequencing in the offspring hypothalamus

Small RNA sequencing was conducted in 8-week-old male offspring of control and obese mothers fed a chow diet (Off-C Chow and Off-Ob Chow groups), to avoid confounding factors of HFD consumption on hypothalamic gene expression. After culling by cervical dislocation, brains were immediately frozen and stored at −80*°*C until sectioning at 12 μm on a cryostat (Leica Biosystems, Wetzlar, DE). PVH and ARC tissue was collected from brain sections using laser capture microdissection with a PALM Robot Microbean (Carl-Zeiss, Gottingen, DE), and RNA extracted as described below. Library preparation and small RNA sequencing of the PVH and ARC samples was performed as described previously (40) using a TruSeq^®^ Small RNA Library Preparation kit (Illumina, Cambridge, UK). Libraries were sequenced using a HiSeq 4,000 platform. Raw reads were mapped to the mouse genome through bowtie v1.2.3. Reads per Kilobase of transcript per Million mapped reads were produced from RNA Sequencing raw output and statistically analysed through likelihood ratio test using edgeR version 3.36.0. A P value cut off of <0.05 was used to determine significantly regulated miRNAs. Sequencing data has been deposited on the GEO data repository.

### 2.3. Cell culture and in vitro treatments

For neural progenitor cell culture, fetal hypothalamus (embryonic day 13) was rapidly dissected in ice cold buffer and neural progenitor cells were harvested by centrifugation (according to (41)) and cultured in Neurocult^TM^ proliferation media (Stem Cell Technologies, Cambridge UK) containing recombinant human EGF according to manufacturer’s instructions. Cells were cultured for 8 days *in vitro* before extraction of RNA as described below. mHypoE-N46 cells (generously donated by Dr. Clemence Blouet) were cultured in growth medium (High glucose Dulbecco’s modified Eagle Medium supplemented with 5% fetal bovine serum, 1% Penicillin-Streptomycin, 2% L-glutamine) and treated for 24h with glucose (20 mM), oleic acid (250 *μ*M), linoleic acid (250 *μ*M), palmitic acid (250 *m*M) or stearic acid (250 *μ*M) (all from Sigma-Aldrich, Saint Louis, Missouri, USA). Mouse brain astrocyte cell line (C8-D1A, CRL-2541, ATCC ®, Manassas, VA USA) were cultured in ATCC-formulated Dulbecco’s Modified Eagle’s Medium, (Catalog No. 30-2002) supplemented with 10% fetal bovine serum and treated for 24h with glucose (20 mM), oleic acid (250 *m*M), palmitic acid (250 *μ*M) or stearic acid (250 *μ*M). After respective treatments, RNA was extracted as described below. To confirm regulation by miR-505-5p of predicted miR targets, mHypoE-N46 cells were transfected with a miR-505-5p mimic (miRCURY LNA miRNA mimic, Qiagen) or negative control, using Lipofectamine-3000 transfection kit (Invitrogen, Waltham, Massachusetts, USA). After 48h of transfection, RNA was extracted for gene expression analyses.

### 2.4. Pulsed stable isotope labelling by amino acids in cell culture (pSILAC)

pSILAC proteomic methodology was performed as described previously (13). Briefly, GT1-7 cells were cultured in regular DMEM media containing light lysine (Lys 0) in two cell culture dishes for 24h and then transfected with miR-505-5p mimic or negative control. After 8h, normal growth medium from both cell populations was replaced with SILAC media containing either heavy lysine [L-lysine-^13^C-^15^N (Lys8)] or medium-heavy lysine [L-Lysine-^2^H (Lys4)] for the miR-505-5p mimic cells and miRNA negative control cells, respectively. After 24h, protein extracts from both cell populations were combined and subjected to SDS-PAGE separation, digested with trypsin and submitted to high performance liquid chromatography tandem mass spectrometry (HPLC/MS/MS). Raw signal data were processed in Maxquant 1.5.2.8 using a Uniprot Mus musculus database (downloaded on 29 Jan 2016, 24,802 sequences). Results were merged with predicted targets for miR-505-5p obtained from Target Scan (http://www.targetscan.org/vert_72/). After bioinformatic analysis, differentially expressed transcripts were validated by qPCR analysis. Pathway enrichment analysis was performed in Ingenuity® Pathway Analysis (Qiagen). pSILAC raw data is available in Supplementary data file 2.

### 2.5. RNA extraction and gene expression analysis

All RNA extraction from tissue and cultured cells was performed using the miRNeasy Micro Kit (Qiagen) to isolate total RNA including small RNAs. cDNA was reverse transcribed using a RevertAid First Strand cDNA Synthesis Kit (Thermo Scientific) or miRCURY LNA RT Kit (Qiagen Sciences) for analysis of mRNA or miRNA respectively. Gene expression was analysed using a StepOnePlus™ Real-Time PCR System (Thermo Fisher Scientific, USA) and either Power SYBR Green PCR Master Mix or Taqman Universal PCR Master Mix (both Thermo Fisher Scientific, USA). Relative quantification of mRNA and miRNA was calculated by the 2^-ΔΔCt^ method. Data were normalized to the geometric average of housekeeping genes *Snord68* and *Unisp6* when miRNA was analysed or to *Sdha* when target genes were analysed (the expression of housekeepers did not change between groups) and values were reported as fold changes compared to values obtained from the control group (set at 1.0). Primers for SYBR Green-based PCR of mRNA were obtained from Sigma Aldrich, miRCURY LNA assays for qPCR of miRNA were purchased from Qiagen, and TaqMan probes were obtained from ThermoFisher Scientific. All primer sequences and assay IDs are listed in Supplementary table 1.

### 2.6. Fatty acid uptake

mHypoE-N46 cells were cultured in a 24 well plate and transfected with miR-505-5p mimic or negative control as described above. After 48h, cells were treated with 1 μM [1-^14^C]-oleic acid (PerkinElmer) diluted in high glucose DMEM containing 0.05% BSA for 1h. Cells used for protein quantification were incubated in uptake medium not containing the radionuclide. After the incubation, media from wells were removed and transferred to 4 ml of scintillation fluid. Cells were carefully washed three times in 1 ml ice cold 0.1% fatty acid free BSA in PBS. After washing, cells were scraped into 200 μl RIPA buffer, transferred to microcentrifuge tubes and centrifuged at 16,000 g for 10 min. 100 μl of lysate was added to 4 ml of scintillation fluid and a beta scintillation counter used to measure concentration of radionuclide. The fatty acid uptake rate was calculated using the concentration of radionuclide, incubation time and total protein amount (and expressed as pmol/min/mg).

### 2.7. Fatty acid metabolism

A fatty acid oxidation protocol was adapted from Pantaleao *et al.* (40). Briefly, mHypoE-N46 cells were cultured in a 24 well cell culture dish and transfected with miR-505-5p mimic or negative control in triplicate. After 48h of transfection, cells were washed with warm PBS and incubated in 500 *μ*l of High-glucose DMEM containing 12.5 mM HEPES, 0.3% fatty acid-free BSA, 1 mM L-carnitine, 100 µM oleic acid and 0.4 μCi/ml [1-^14^C]-oleic acid. 200 *m*l of concentrated hydrochloric acid was injected into the wells, and the acid solution was left to incubate for an extra hour. The acid solution was then centrifuged at maximum speed (≥ 14,000 × g) for 10 min at 4 °C and 500 µl supernatant transferred to a scintillation vial for radioactivity measurement of acid soluble metabolites from fatty acid metabolism (Acetyl-CoA, acyl-carnitines etc).

### 2.8. Characterisation of miR-505-5p targets in human datasets

miR-505-5p fatty acid metabolism related target genes were mapped to orthologous genes in the human genome using ENSEMBL and integrated with common variant genome-wide association study (GWAS) data for body mass index (BMI), from the GIANT BMI study (42). Each of the miR-505-5p target genes was annotated based on proximity to genome-wide significant signals (p<5×10^-8^), in 1Mb windows; 500kb up- or downstream of the gene’s start or end site. For genes with proximal GWAS signals, we calculated genomic windows of high linkage disequilibrium (LD; R^2^>0.8) for each given signal and mapped these to the locations of known enhancers for the target genes, using the activity-by-contact (ABC) enhancer maps (43). We also performed a gene-level Multi-marker Analysis of GenoMic Annotation (MAGMA) analysis (44), which collapses all observed genomic variants within a given gene and calculates aggregate gene-level associations to the phenotypic trait. Genes exhibiting an FDR-corrected MAGMA p-value <0.05 were considered significant. Finally, for genes in loci containing at least a suggestive level of GWAS significance (p<5×10^-5^), we performed summary-data-based Mendelian randomization (SMR) and heterogeneity in dependent instruments (HEIDI) tests(v1.02; (45)) using blood expression quantitative trait loci (eQTL) data from the eQTLGen study (46). For the eQTL analyses, we considered gene expression to be associated with BMI if the FDR-corrected p-value for the SMR-test was p<0.05 and the p-value for the HEIDI test was >0.001.

### 2.9. Intra-cerebroventricular (ICV) injection of miR-505-5p mimic

Adult C57Bl/6J male mice were implanted with an intra-cerebroventricular (ICV) canula (Plastics One, Inc.) in the lateral ventricle (coordinates from Bregma: anteroposterior −0.4; mid-lateral −1.0 and dorsoventral −2.3 mm). One week prior to the ICV surgery, mice were exposed to a high fat diet (60% fat) for familiarization and for basal food intake measurement. After 5 days of recovery from surgery, mice received a 0.5μM dose of a miR-505-5p mimic (Qiagen) or negative control combined with HiPerfect transfection reagent (Qiagen) into the lateral ventricle. The ICV treatment was repeated once every 24h for 3 days through the cannula and *ad lib* food intake was measured once every 24h (as shown in Figure 5a).

### 2.10. Statistics

Data were analysed using Prism 8 (Graphpad, La Jolla, USA) by different tests according to the numbers of groups and time points: unpaired or paired t-test (as specified) for experiments with two treatment groups, one-way ANOVA for experiments with 3 or more groups at one time point, mixed effects analysis for experiments with 2 or more groups over time. Tukey or Sidaks multiple comparisons test were performed after mixed effects analysis (as specified in figure legends). Outliers were detected by Grubb’s method. For pSILAC, statistical analysis of normalized labelled: unlabelled protein ratios was performed using multiple linear models with Bayesian correction using limma package (v.2.16.0) for R (v.4.1.2). All data are shown as mean *±* SEM. In each case n refers to the number of different litters represented.

### 2.11. Study approval

The animal studies performed in this manuscript were licensed by the UK Home Office under project license P5FDF0206 and approved by the University of Cambridge Animal Welfare and Ethical Review Body and by the Ethics Committee of Animal Welfare from Instituto de Ciencias Biomedicas-Universidade de Sao Paulo (protocol number 72/2017).

## 3. Results

### 3.1. Male offspring born to obese mothers show increased intake of a high-fat diet and subsequent increased adiposity

Maternal diet-induced obesity during pregnancy has been suggested to program increased food intake and subsequently adiposity in the offspring. We recorded food intake, body weight and adiposity of male offspring born to obese and control mothers, who were weaned onto either a normal chow or an obesogenic high fat diet (HFD) from weaning until 8 weeks of age (Figure 1a). Male offspring of obese mothers (Off-Ob chow) did not consume more calories during *ad lib* feeding over this period when weaned onto a chow diet compared to the offspring of lean mothers fed a chow diet (Off-C chow) (Figure 1b). However, when weaned onto an HFD, offspring born to obese mothers (Off-Ob HFD) showed significantly higher daily caloric intake from 6 weeks of age compared to offspring of lean mothers weaned onto the same HFD (Off-C HFD) (Figure 1c). In a two-bottle choice test for sucrose performed in male mice, all mice preferred sucrose solution to water (i.e., a preference > 50%) and both sucrose preference and absolute intake of sucrose increased with increasing sucrose concentration (Figure 1d and 1e). There was no significant effect of perinatal exposure to maternal obesity on absolute sucrose preference (Figure 1d) or sucrose intake (Figure 1e) in offspring.

There was no effect of perinatal exposure to maternal obesity on offspring body weight over the period studied (i.e. up to 8 weeks of age), regardless of whether offspring were weaned onto a chow diet (Figure 1f) or a HFD (Figure 1h). Longitudinal adiposity measurements showed increased adiposity in all animals weaned onto a HFD compared to chow fed animals; but this increase in adiposity was greater in offspring of obese mothers (Figure 1i). We have previously shown that energy expenditure is unchanged in male mice exposed to maternal obesity at 8 weeks of age (14) so the increased intake of a HFD is the likely cause of increased adiposity.

### 3.2. Maternal obesity programs increased expression of miR-505-5p in offspring hypothalamus from fetal to adult life

In order to investigate the effect of exposure to maternal obesity on miRNA expression profile in the offspring hypothalamus, we analysed the miRNA profile of two key nuclei, the paraventricular nucleus (PVH) and arcuate nucleus (ARC) of the hypothalamus, in young adult chow-fed male mice born to obese mothers or lean control mothers. Small RNA sequencing analysis highlighted 4 miRNA that were dysregulated in the PVH (Suppl. Table 2) and 14 in the ARC (Suppl. Table 3) of offspring exposed to maternal obesity. The miRNA miR-505-5p, which is conserved between mice and humans (Figure 2a) showed a striking increased expression in both the ARC (Figure 2b) and PVH (Figure 2c) of offspring born to obese mothers in comparison to offspring of control mothers. The results of the small RNA sequencing analysis were validated by qPCR for miR-505-5p in a separate cohort of young adult male mice (Fig 2d). In control animals, miR-505-5p was expressed at a higher level in the hypothalamus than two other regions of the brain we examined (cortex: 1.00*±* 0.06, hippocampus: 1.64 *±* 0.14, hypothalamus: 2.66*±* 0.19 (A.U. normalised to cortex)). In contrast to findings in the hypothalamus, exposure to maternal obesity did not result in altered expression of miR-505-5p in either the cortex (OffC: 1.00 *±* 0.06, OffOb: 1.01 *±* 0.10 (A.U. normalised to OffC cortex)) or hippocampus (OffC: 1.64 *±* 0.14, OffOb: 1.41 *±* 0.36 (A.U. normalised to OffC cortex)), confirming that exposure to maternal obesity increases expression of miR-505-5p specifically in the hypothalamus.

**Figure 2.**
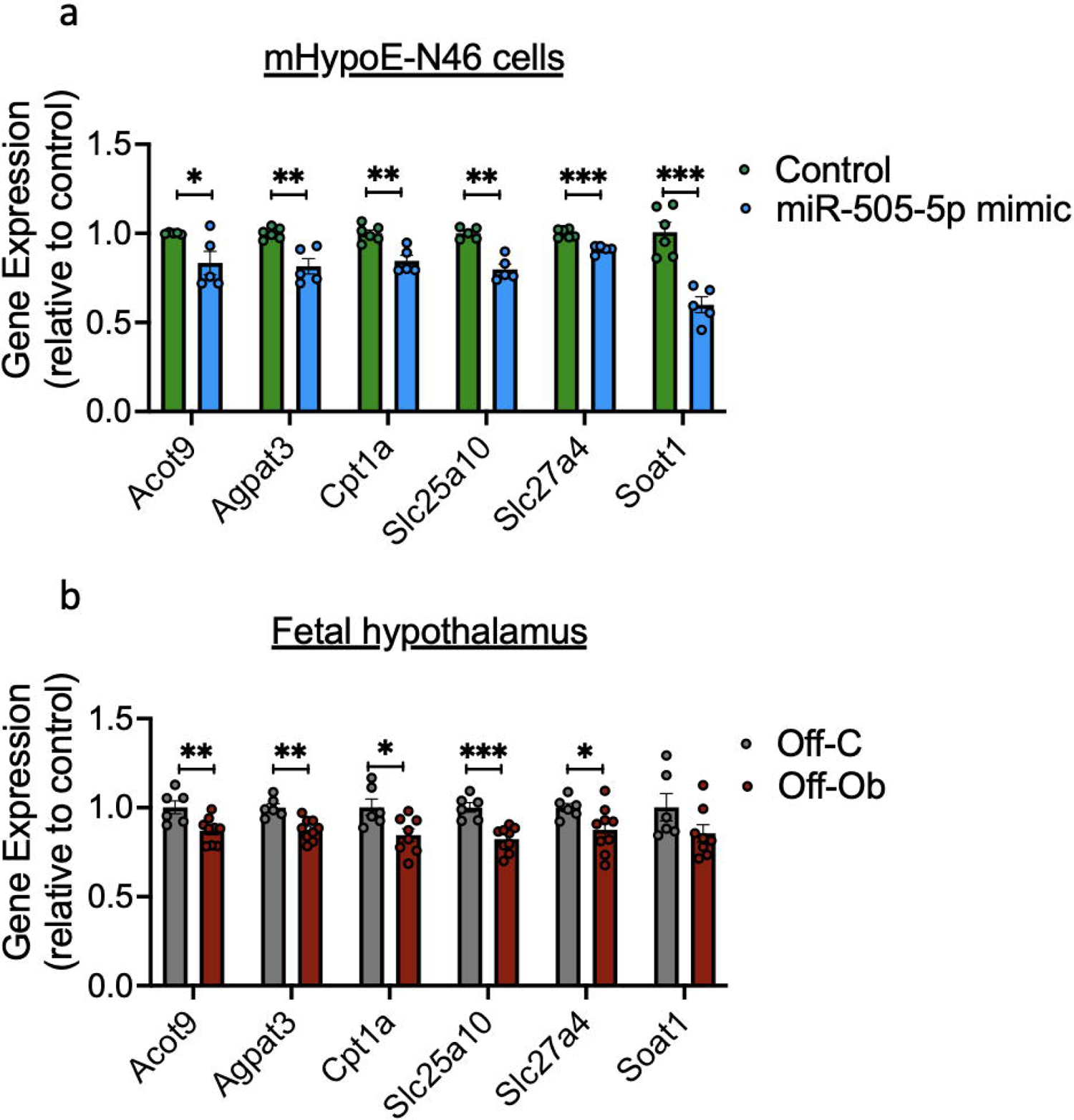
Hypothalamic miR-505-5p expression is increased in offspring of obese mothers from fetal to adult life, and *in vitro* in neurons by exposure to metabolic hormones and fatty acids. a) Sequence overlap of mature miR-505-5p in mice and humans. b & c) Volcano plot of small RNA sequencing results showing significant overexpression of miR-505-5p (red dot surrounded by blue diamond) in the ARC (b) and PVH (c) of 8-week-old offspring born to obese mothers compared to offspring of lean mothers. d-f) qPCR analysis of miR-505-5p expression in (d) ARC at 8 weeks of age; (e) in whole hypothalamus at E13; (f) in hypothalamic neural progenitor cells extracted at E13 and cultivated as neurospheres for 8 days *in vitro*. g) *Atp11c* (miR-505-5p host gene) mRNA levels in E13 whole hypothalamus. h) miR-505-5p relative expression of mHypoE-N46 cells exposed to insulin or leptin. i) miR-505-5p relative expression of mHypoE-N46 cells exposed to oleic acid, palmitic acid, stearic acid, linoleic acid and glucose for 24h. j) miR-505-5p relative expression of C8-D1A (astrocyte) cells exposed to oleic acid, palmitic acid, stearic acid and glucose for 24h. k) miR-505-5p relative expression in mediobasal hypothalamus (MBH) in lean, diet-induced obese or genetically obese (Ob/Ob) male mice.* P<0.05, **P<0.01. Statistical significance was determined with unpaired t-test. Outliers were excluded from (g) for obese group (outlier excluded value 0.878) and j for oleic acid treatment (outlier excluded value 1.554). The underlying data are provided as a Source Data file

In order to ascertain how early in life the over-expression of miR-505-5p is established, we analysed miR-505-5p expression in fetal hypothalamic samples and found a significant upregulation of the miRNA in E13 fetuses exposed to maternal obesity (Figure 2e). To determine if this overexpression is maintained outside the intrauterine environment, which would suggest a cell-autonomous programmed effect, we analysed hypothalamic NPC from E13 offspring and found overexpression of miR-505-5p in NPC after 8 days grown *in vitro* (Figure 2f). miR-505-5p sits in the gene *Atp11c*, a type IV P-type ATPase. The expression of *Atp11c* was also increased (though to a lesser extent than the miRNA) in the hypothalamus of fetuses exposed to maternal obesity, showing that miR-505-5p is at least in part regulated along with the host gene (Figure 2g). These data show that miR-505-5p overexpression is permanently programmed in the hypothalamus following perinatal exposure to maternal obesity and is a potential candidate to mediate the long-term effects of maternal overnutrition on offspring hypothalamic function.

### 3.3. Insulin and specific fatty acids increase the expression of miR-505-5p in a hypothalamic neuronal cell line *in vitro*

We next sought to uncover which factors related to maternal obesity could be responsible for the increase in miR-505-5p in hypothalamus. We have previously reported an increase in circulating levels of insulin and leptin in the serum of diet-induced obese dams in our mouse model (37). Application of insulin, but not leptin, to a hypothalamic neuronal cell line (mHypoE-N46) caused an increase in expression of miR-505-5p (Fig 2h). We have previously conducted mass spectrometry analysis of the HFD pellet to establish the primary FA components of the obese dam diet, and found the FAs in greatest abundance were palmitic, stearic, oleic and linoleic acid (47). We therefore sought to identify whether any of these FAs can alter the expression of miR-505-5p in mHypoE-N46 cells. The expression of miR-505-5p was increased in neurons after oleic and stearic acid treatment (Figure 2i). Neither palmitic nor linoleic acid treatment caused overexpression of miR-505-5p in neurons (Figure 2i). As the obesogenic diet fed to obese dams is supplemented with condensed milk which is high in sucrose, we also tested to see whether glucose could regulate expression of miR-505-5p *in vitro*. Glucose did not alter the expression of miR-505-5p in hypothalamic cells *in vitro* (Figure 2i). As astrocytes are also reported to be involved in hypothalamic FA sensing, we also tested whether miR-505-5p was expressed in C8-D1A cells and if so whether the expression was regulated by components of the obesogenic diet used in our model. Although miR-505-5p is expressed in astrocytes, it was not regulated by FAs in the same manner we observed in neurons (Fig 2j). In fact, we observed a down-regulation of miR-505-5p expression in response to palmitic acid in astrocytes (Fig 2j).

In addition to regulation by nutrients and hormones known to be altered in pregnancies complicated by obesity, we also examined whether miR-505-5p levels are modulated *in vivo* by long-term obesity. We used two separate obesity models to delineate the effects of obesity *per se* from consumption of a high-calorie diet: diet-induced obese animals and leptin deficient *Ob/Ob* animals (fed a chow diet). The expression of miR-505-5p in the hypothalamus was increased in male mice with diet-induced obesity (Figure 2k) and there was a trend towards increase in mice with genetic obesity (analysed by one way ANOVA). This shows that hypothalamic miR-505-5p is regulated by current obesity in adult animals (independent of an HFD), as well as by factors associated with maternal obesity.

### 3.4. miR-505-5p targets identified by pSILAC are involved in FA uptake and metabolism

miRNAs are known to regulate gene function through degradation of mRNAs and/or via inhibition of protein translation. We performed a proteomic analysis to identify targets of miR-505-5p by pSILAC following transfection of a miR-505-5p mimic in a hypothalamic cell line, GT1-7 (Figure 3a). Cumulative fraction analysis showed that proteins encoded from mRNAs containing binding sites for miR-505-5p according to TargetScan (v. 7.2) were downregulated by the miR-505-5p mimic, showing that the pSILAC was effective in identifying proteins regulated as a consequence of over-expression of miR-505-5p (Figure 3b). pSILAC data was overlaid with predicted targets containing the miR-505-5p seed sequence identified by TargetScan and miRbase to identify potential direct targets (Figure 3c).

**Figure 3.**
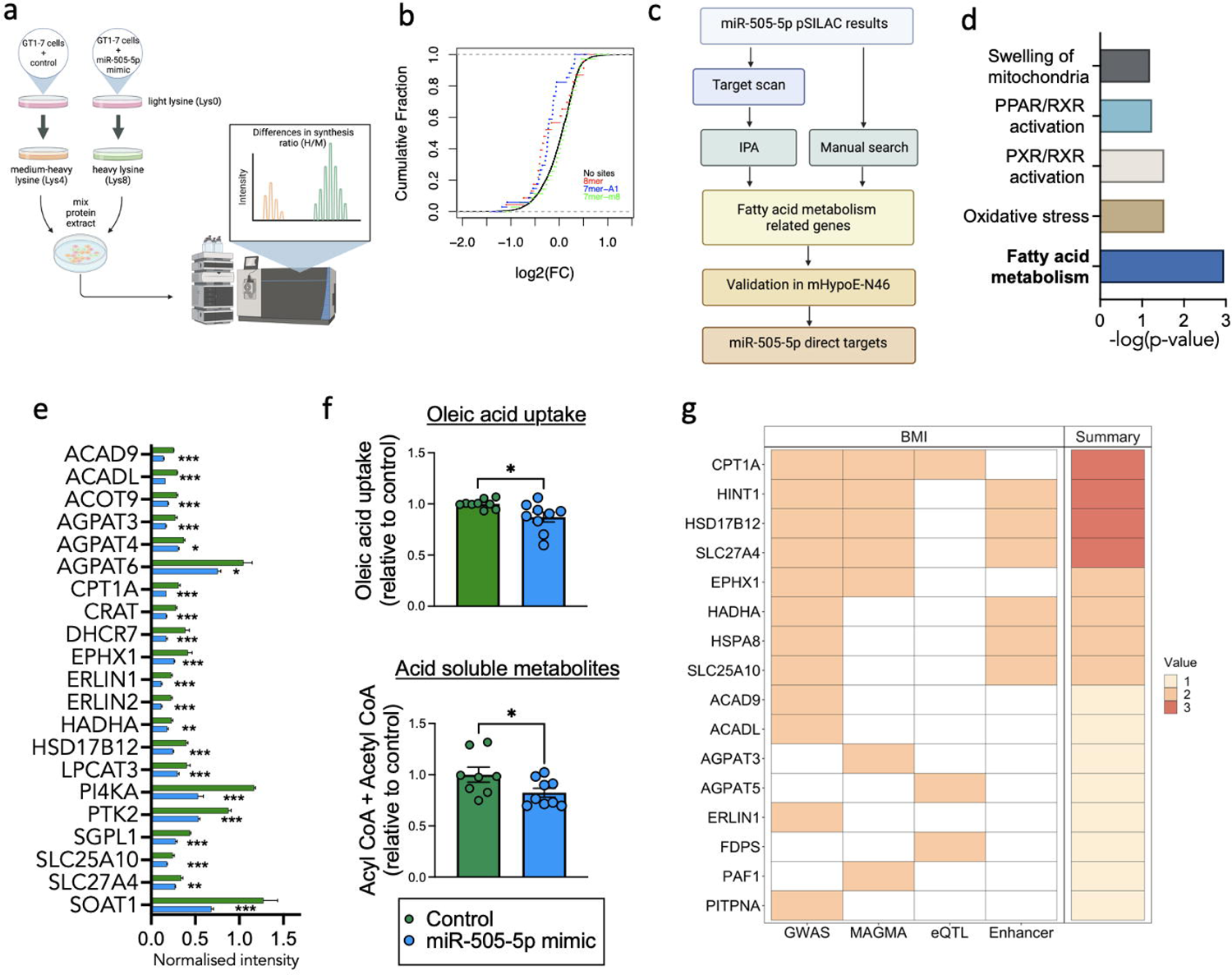
miR-505-5p targets are involved in fatty acid metabolism and genetically associated with BMI in humans. **a)** Diagram of pulsed SILAC experiment in GT1-7 cells to identify targets of miR-505-5p b) Cumulative fraction analysis of lys8-labelled: lys4-labelled proteins ratio according to occurrence and type of mRNA binding site for miR-505-5p, as predicted by TargetScan (v. 7.2) c) Workflow used to identify direct targets of miR-505-5p. d) Top regulated pathways among miR-505-50 direct targets as reported in Ingenuity Pathway Analysis (IPA). e) miR-505-5p target proteins of interest identified through IPA and manual search of pSILAC data. f) Radiolabelled oleic acid metabolism into acid-soluble metabolites and oleic acid uptake in mHypoE-N46 following transfection with either a miR-505-5p mimic or control for 48h. g) Heatmap showing the overlap between human orthologs of identified miR-505-5p fatty acid metabolism related target genes in mice and human genetic datasets. Target genes were annotated on the basis of (i) proximity to GWAS signals, (ii) aggregate gene-level associations to the trait, (iii) colocalization between the GWAS and eQTL data, and (iv) the presence of known enhancers within the GWAS signals. Total number of the observed concordant predictors (out of 4) is displayed in the summary panel. Expanded results can be found in Supplementary Table 4. *p<0.05, ** p<0.01, *** p<0.001. Statistical significance was determined by unpaired T-test. The underlying data are provided as a Source Data file

Ingenuity Pathway Analysis (IPA) revealed that miR-505-5p targets are enriched for processes related to fatty acid metabolism, followed by oxidative stress and mitochondrial processes (Figure 3d). After conducting a manual search in addition to the IPA, we identified 30 proteins related to FA metabolism pathways that were significantly dysregulated in the pSILAC experiment. Twenty-one of these FA related proteins were downregulated after miR-505-5p over-expression (Figure 3e), suggesting an overall reduction in FA metabolism processes in the cells. We next investigated the functional consequences of over-expression of miR-505-5p in a hypothalamic cell line on FA oxidation and FA uptake processes. We used radio-labelled oleic acid to show that oleic acid metabolism into acid-soluble metabolites (Figure 3f) and oleic acid uptake (Figure 3f) were decreased in hypothalamic neurons after miR-505-5p over-expression. As FA uptake and metabolism to Acetyl-CoA and TCA cycle intermediates are thought to be key steps through which neurons are able to sense circulating FA levels, these results suggest that increased miR-505-5p *in vivo-* through an impact on hypothalamic fatty acid uptake and oxidation-could alter the subsequent ability of the neurons to sense circulating LCFA levels.

### 3.5. miR-505-5p and its targets are associated with BMI in humans

A recent study of 3000 adults has identified circulating miR-505-5p levels as being one of only a few miRNA positively associated with BMI in humans (48). To further understand the contribution of miR-505-5p targets to human metabolic health, we mined phenotypic and genetic data from human population studies by assessing data from a common variant GWAS of BMI (N=806,834)(42). Of the 30 human orthologues to the FA metabolism related targets of miR-505-5p, 12 genes were proximal to common variant genome-wide significant signals; when variants were analysed at the gene-level using MAGMA, 7 genes were significantly associated with BMI (Suppl. Table 4). As most common variant GWAS signals are intronic or intergenic, we overlayed these associations with other datasets to understand whether the associated variants can be functionally linked to our target genes (Figure 3g). Genomic variants within 6 of the BMI association signals are within regions of known enhancers(43). We also observed colocalization between these signals and gene expression levels for 3 target genes.

Across these analyses, our human genetic analyses had the greatest support for CPT1A, HINT1, HSD17B12 and SLC27A4, which had the greatest number of variant to gene predictors (Figure 3g). For CPT1A, the BMI GWAS colocalised with CPT1A eQTLs (FDR-corrected P SMR=5.94×10^-5^ and P HEIDI=0.002, Suppl. Table 4), while signals proximal to HINT1, HSD17B12 and SLC27A4 all overlapped with known enhancers for these genes. Collectively, these observations suggest that several of the novel miR-505-5p target genes we identified are functionally associated with increased BMI in humans.

### 3.6. miR-505-5p overexpression in vitro, as well as in utero exposure to maternal obesity, causes downregulation of genes related to FA sensing in hypothalamic cells

As described above, a manual search combined with the IPA identified 30 genes linked to FA metabolism (Figure 3e). Among the genes related to FA metabolism that were downregulated by miR-505-5p were genes related to fatty acid transport (*Slc25a10* and *Slc27a4* (that encodes the protein FATP4)) and fatty acid oxidation (*Acot9*, *Agpat3*, *Cpt1a* and *Soat1*). To confirm the downregulation of predicted miR-505-5p targets relevant to FA metabolism, we measured expression of these genes following transfection of a miR-505-5p mimic in the mHypoE-N46 hypothalamic cell line. miR-505-5p mimic treatment decreased the mRNA expression of *Acot9*, *Agpat3*, *Cpt1a*, *Slc25a10, Slc27a4* and *Soat1* (Figure 4a), compared to cells treated with a vehicle control.

**Figure 4.**
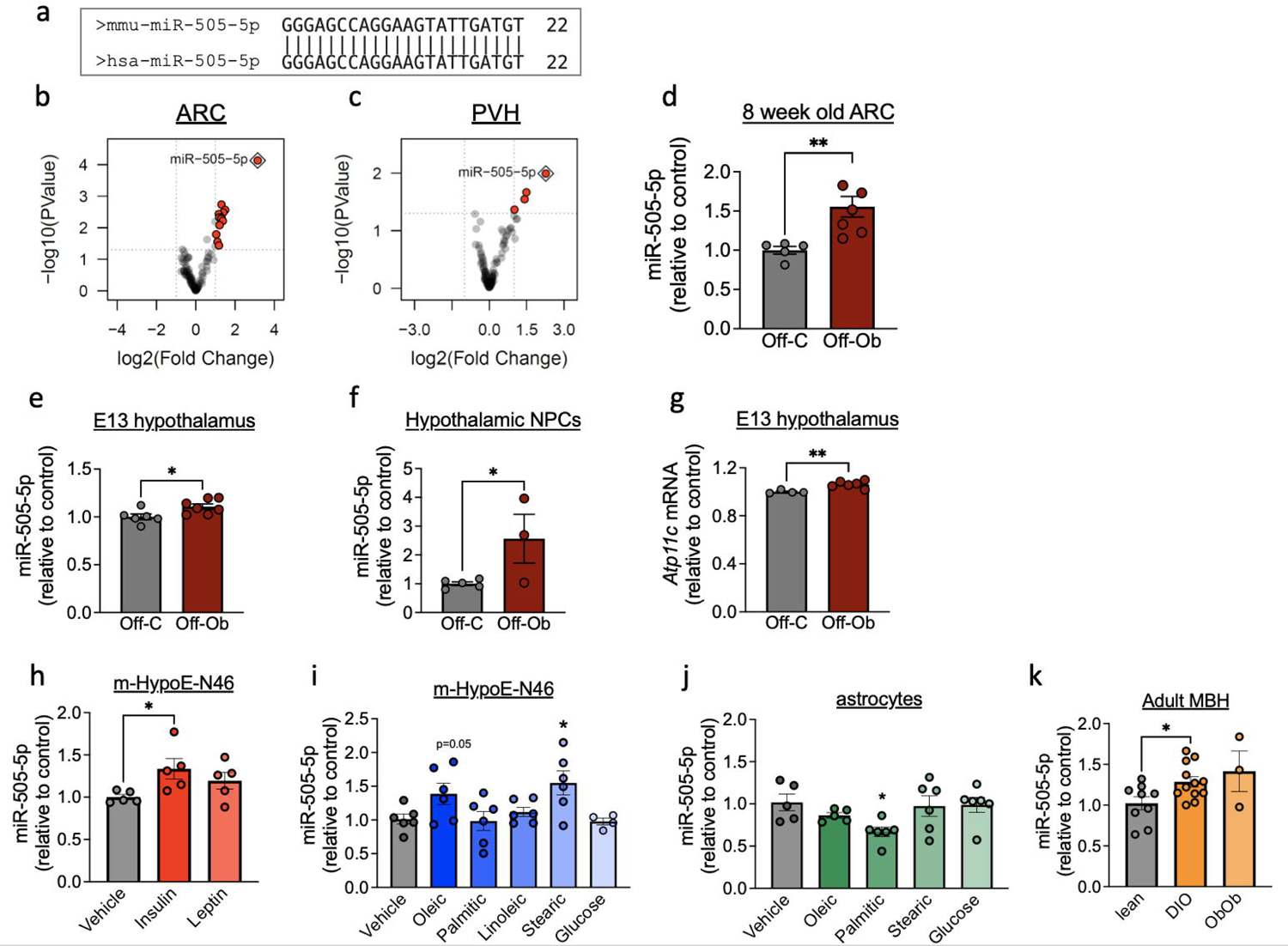
miR-505-5p target genes related to fatty acid sensing are downregulated after treatment with miR-505-5p *in vitro* and in fetal hypothalamus of offspring from obese mothers. **a)** Relative mRNA expression of genes related to lipid metabolism in mHypoE-N46 cells after treatment with a miR-505-5p mimic and (b) in the hypothalamus of fetuses from control or obese pregnancy at E13 * P<0.05, ** P<0.01, *** P<0.001. Statistical significance was determined by unpaired T-test. Outliers were excluded from (a) for miR-505-5p mimic (Cpt1a outlier excluded value 1.19, Soat1 outlier excluded value 1.71). The underlying data are provided as a Source Data file

**Figure 5.**
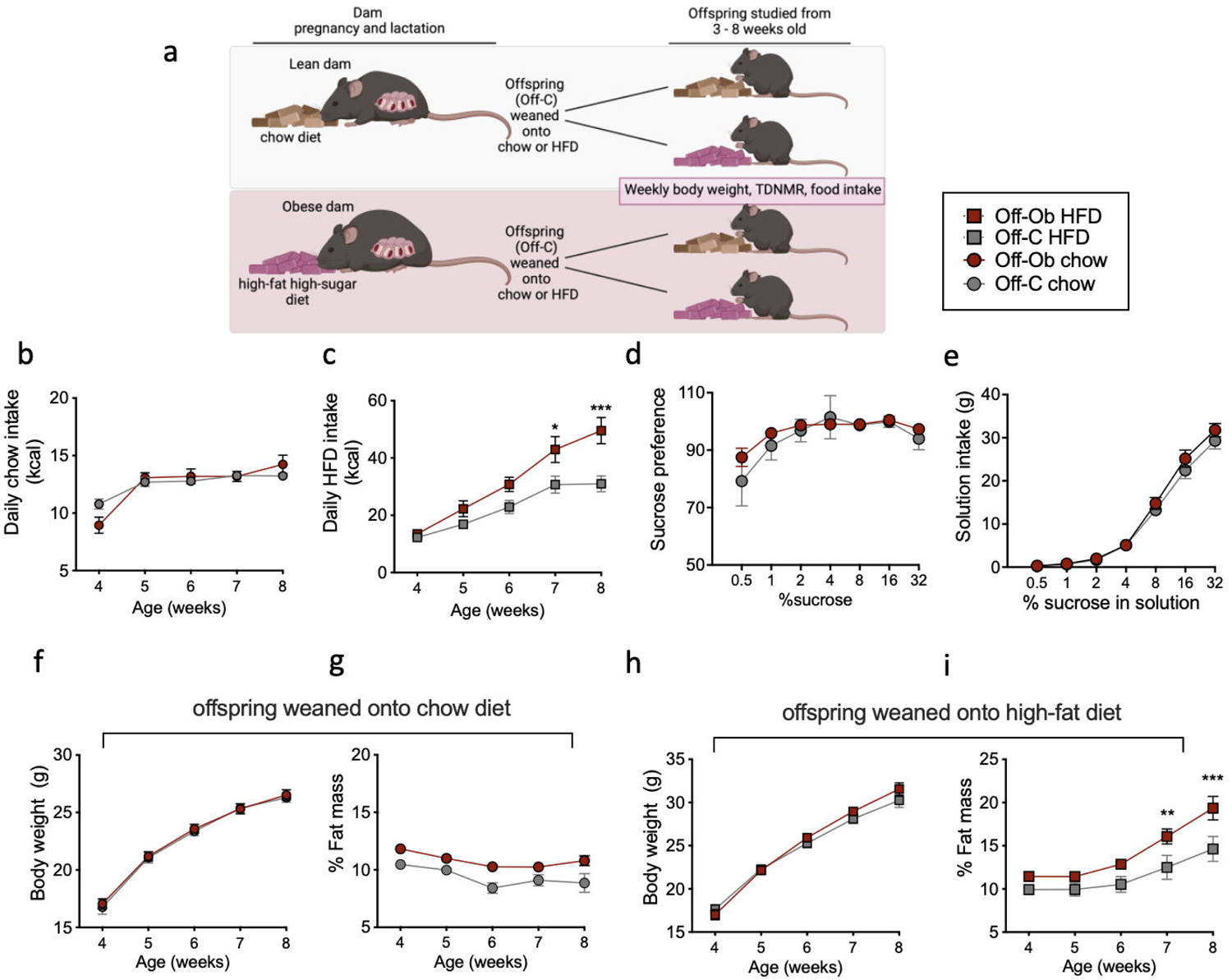
Intra-cerebroventricular injection of miR-505-5p in adult mice increases intake of a high-fat diet, whilst a maternal exercise intervention corrects offspring hypothalamic miR-505-5p levels and reduces intake of a high-fat diet. a) Diagram depicting surgical schedule. Mice were allowed to recover for 5 days post-implanted of an ICV cannula into the lateral ventricle. Mice then received injection of either a vehicle or miR-505-5p mimic into the ICV cannula and food intake was measured over 3 days. b) qPCR confirming over-expression of miR-505-5p in the hypothalamus after ICV injection of miR-505-5p mimic. c) Chow diet cumulative food intake after 3 consecutive days of treatment (24-1 dose, 48-2 doses and 72 hours-3 doses-after first treatment). d) High-fat diet cumulative food intake after 3 consecutive days of treatment (24-1 dose, 48-2 doses and 72 hours-3 doses-after first treatment). e) Graphical depiction of maternal obesity model with third group of dams that underwent moderate exercise intervention during obese pregnancy. f) relative expression of miR-505-5p in the arcuate nucleus of 8 week old male offspring born to control, obese or obese-exercised dams. g) HFD pellet daily kilocalorie intake of male offspring born to control, obese or obese-exercised dams * p<0.05, ** p<0.01. Statistical significance was determined by unpaired T-test (c & d), one way ANOVA (f), and mixed effects analysis with Sidak post-hoc test (g). The underlying data are provided as a Source Data file.

Decreased mRNA expression of the genes *Acot9*, *Agpat3*, *Cpt1a*, *Slc25a10* and *Slc27a4* was also observed in the fetal hypothalamus at E13 of offspring from obese mothers, compared to the offspring of lean mothers (Figure 4b). This is consistent with our observed increase in hypothalamic miR-505-5p levels at E13 (Figure 2e). These data show that both over expression of miR-505-5p in a hypothalamic cell line mHypoE-N46 *in vitro*, and exposure to maternal obesity *in vivo* - which we have shown programs increased miR505-5p in the hypothalamus - downregulate the expression of genes related to FA metabolism.

### 3.7. Intra-cerebroventricular injection of miR-505-5p increases high-fat-but not chow-pellet intake

Our results demonstrate that targets of miR-505-5p in the hypothalamus include genes involved in neuronal FA sensing, and that manipulation of miR-505-5p levels reduces rates of neuronal FA uptake and metabolism. Previous studies have suggested that neuronal FA uptake and metabolism are key processes that allow neurons to sense circulating FA levels and initiate appropriate adjustments to parameters including food intake (22). We therefore hypothesised that over-expression of miR-505-5p would alter the ability of hypothalamic neurons to sense circulating FA levels and result in a change in food intake. To verify whether miR-505-5p has a direct effect *in vivo* in the brain in FA sensing - and therefore food intake - we performed ICV administration of a miR-505-5p mimic or a negative control and measured *ad lib* intake of high-fat diet or chow diet for 3 consecutive days following surgery (Figure 5a). Increased miR-505-5p in the hypothalamus after ICV injection of the miR-505-5p mimic was confirmed on post-mortem dissection (Figure 5b). Mice that received the miR-505-5p mimic ICV had increased high-fat diet intake compared to mice that received a negative control treatment (Figure 5c). The increase in food intake caused by ICV miR-505-5p was exclusively seen when animals were given the high fat diet; ICV miR-505-5p did not increase food intake when animals were fed a chow diet (Figure 5d).

### 3.8. Maternal exercise intervention in an obese pregnancy rescues the programmed over-expression of hypothalamic miR-505-5p and normalises high-fat diet intake in the offspring

We have previously shown that a moderate exercise intervention in obese dams during pregnancy rescues maternal insulin sensitivity-independent of changes in adiposity (39)- and is associated with a rescue of the cardiac dysfunction observed in offspring exposed to maternal obesity (49) (Figure 5e). We therefore sought to investigate whether the correction of maternal metabolic health was sufficient to rescue the programmed over-expression of hypothalamic miR-505-5p offspring and changes in offspring feeding behaviour. Offspring of obese exercised dams (Off-FEx) show a reduced expression of miR-505-5p in the ARC compared to Off-Ob to a level comparable to that observed in controls (Figure 5f). As we have shown that over-expression of miR-505-5p in the hypothalamus causes increased intake of a high-fat diet, we would expect Off-FEx to show a reduced intake of HFD compared to Off-Ob. Indeed, offspring of obese-exercised dams with normalised expression of miR-505-5p show significantly reduced intake of a HFD from 7 weeks of age, despite their exposure to perinatal obesity (Figure 5g). At this age there is no difference in body weight between the three groups of offspring (OffC 25.65g *±* 0.67, OffOb 26.66g *±* 0.32, OffFEx 25.67g *±*0.63). Therefore effects of the maternal exercise intervention are not related to changes in offspring body weight.

## 4. Discussion

In the current study, we have used a rodent model to examine the effects of exposure to maternal obesity on offspring food intake, small RNA sequencing in the hypothalamus to identify programmed miRNAs and a series of *in vitro* and *in vivo* experiments-including human genetic analysis-to establish how miRNA may causally influence feeding behaviour. Our results showed that maternal obesity alters the miRNA profile in the ARC and PVH and identified miR-505-5p as a miRNA that is programmed in the fetal hypothalamus with changed expression maintained through to adulthood. We have shown that targets of miR-505-5p are related to FA uptake and metabolism, that some of these targets are associated with increased BMI in humans, and that manipulation of miR-505-5p levels in hypothalamic neurons *in vitro* alters these processes. Importantly, we have shown that ICV injection of a miR-505-5p mimic increases intake specifically of an HFD, and an exercise intervention in obese dams rescues programmed miR-505-5p levels and reduces HFD intake. Altered expression of hypothalamic miR-505-5p *in utero* is therefore a novel route by which perinatal exposure to maternal obesity programs permanent changes to offspring feeding behaviour, leading to obesity.

The increased intake of an HFD in the offspring of obese mothers precedes the increased adiposity observed in these animals and given that we have previously reported no differences in energy expenditure in mouse offspring exposed to maternal obesity, suggests a causal relationship. Indeed, many studies have now linked changes in food intake to the increased susceptibility to develop obesity in animals exposed to obesity in the perinatal period, and suggested that altered hypothalamic development and/or function plays a key role in this process (4). To our knowledge, we are the first to conduct small RNA sequencing to identify the full spectrum of miRNAs that are dysregulated in the ARC and PVH following exposure to maternal obesity. A role for hypothalamic miRNAs in maintaining energy homeostasis is largely unexplored, but deletion of the miRNA processing enzyme DICER in POMC neurons results in obesity (50), and hypothalamic miR-29a and miR-103 have been shown to protect against obesity (18, 51). Although a few studies have demonstrated altered hypothalamic miRNA expression in mice in response to nutritional status or leptin administration (16, 52, 53), whether exposure to altered nutritional status *in utero* programs hypothalamic miRNA profile was previously unknown. Our small RNA sequencing analysis identified miR-505-5p as being highly up-regulated in both the ARC and PVH of male adult offspring after exposure to maternal obesity when offspring themselves were fed a chow diet and at a time when there was no difference in offspring body weight. We subsequently demonstrated that increased expression of miR-505-5p is detectable as early as E13 in the fetal hypothalamus. Notably, a recent study found miR-505-5p to be one of four miRNAs associated with several markers of cardiometabolic phenotypes in humans (48). The increased expression of miR-505-5p was maintained *in vitro* in hypothalamic neural progenitor cells after one week, indicating this is a programmed effect in the cell, rather than an immediate effect of the intra-uterine environment on gene expression. Interestingly, the miR-505-5p host gene Atp11c, which encodes a Type IV P-type ATPase, a flippase that is involved in the transport of phospholipids, is also increased in the hypothalamus of fetuses of obese mothers. It is intriguing that this apparent nutrient-sensitive miRNA sits within another gene linked to cellular energy levels and that this gene is altered by exposure to maternal obesity. This study does not address the expression or regulation of miR-505-5p in female offspring exposed to maternal obesity and this is an important avenue for future work. Circulating levels of miR-505-5p have been reported to be lower in human females (54), but the Framingham study that reported an association between circulating miR-505-5p and cardiometabolic phenotypes (48) was conducted in a mixed sex group, suggesting miR-505-5p may also play a role in metabolism in females.

The fact that hypothalamic miR-505-5p was up-regulated in both diet-induced obese animals and leptin deficient *Ob/Ob* obese animals shows that consumption of a HFD is not required for miR-505-5p over-expression. This suggests it is a factor related to the maternal obesity *per se* (such as a circulating hormone) that programs over-expression, rather than a metabolite of the HFD consumed by the dam that passes directly to the fetus. Furthermore, this indicates that direct regulation of this miRNA by obesity in adulthood may further compound the ability of an individual with obesity to regulate their food intake. We show that miR-505-5p is increased in response to insulin, oleic and stearic acid in a hypothalamic cell line *in vitro*. We have shown that fatty acids and insulin are increased in the serum of dams fed an obesogenic diet near the end of pregnancy (14), and insulin is also high in fetuses of a human obese pregnancy as well as in our mouse model. Regulation of miR-505-5p by glucose (another metabolic signal known to be high in fetuses exposed to maternal obesity) was ruled out by our *in vitro* data. Interestingly, the human Framingham study that identified miR-505-5p as positively associated with cardiometabolic parameters found no correlation with circulating glucose, and the strongest association was with circulating triglycerides (48) supporting our *in vitro* data. The maternal exercise intervention normalises dam insulin sensitivity, but not maternal body weight, adiposity or leptin levels (39), and this was sufficient to rescue the programmed over-expression of hypothalamic miR-505-5p in offspring exposed to maternal obesity and prevented programmed increased in intake of a high fat diet. Our results therefore suggest that increased circulating free-fatty acids or insulin levels may be key mechanisms by which maternal obesity programs changes to the developing offspring hypothalamus.

As offspring remained in the cage with the dam (who was consuming the obesogenic diet throughout both the gestation and weaning period), we cannot discount the possibility that some of the effects we see on offspring phenotype are due to the juvenile offspring eating the HFD pellets themselves in the latter part of the weaning period. However, it should be noted that exercise intervention during the pregnancy period alone prevented the programmed increase miR-505-5p expression as well as increased food intake. This therefore suggests that targeting the pregnancy period alone is sufficient to prevent at least some of the programmed phenotypes. This issue could be further explored in the future with a cross-fostering experiment where offspring of obese dams could be raised by lean dams consuming a chow diet.

Unlike the well characterised populations of hypothalamic glucose sensing neurons (25), the presence, location and molecular signature of FA sensing neurons in the hypothalamus is not well defined. FAs are able to cross the blood brain barrier (55, 56), and the presence of FA transporters such as FATP1, FATP4 and FAT/CD36, as well as genes required for metabolism of FA (27) support the theory that at least some hypothalamic neuronal populations are FA-sensing. A recent study has shown that knockdown of Free Fatty Acid Receptor-1 in POMC neurons promotes hyperphagia and body weight gain (57). Therefore, it is very likely that FA sensing is a mechanism by which the hypothalamus responds to nutrient status changes, particularly that of an energy surfeit (58). Current hypotheses suggest neuronal cells use the rate of uptake and/or metabolism of FAs to sense circulating levels, (although there may also be direct regulation of some neurons via electrophysiological effects (31, 59) as is the case for glucose-sensing neurons). For this reason, it is relevant that we have shown that miR-505-5p targets include both FA transporters (FATP4) and key components of FA oxidation pathways (such as CPT1A, which has recently been shown to be reduced in offspring in a different model of maternal HFD consumption (60)), and that over-expression of miR-505-5p in a hypothalamic cell line disrupted FA uptake and metabolism. Therefore, altered expression of miR-505-5p caused by *in utero* exposure to high FA levels has the potential to permanently disrupt hypothalamic FA sensing mechanisms. This is supported by the observation that CNS administration of a miR-505-5p mimic results in increased intake of a high-fat diet (but not a chow diet), and that reduced levels of miR-505-5p in offspring of obese exercised dams results in reduced intake of a high-fat diet. Thus, our results support the hypothesis that if neurons cannot sense circulating FA levels due to altered FA uptake and metabolism pathways, animals cannot maintain energy homeostasis, resulting in increased consumption of foods high in fat. This is also supported by a previous study showing that maternal HFD consumption during pregnancy reduced the sensitivity of offspring ventromedial hypothalamic neurons to oleic acid (35).

Although we have highlighted only six critical genes in our experiments, the pSILAC analysis identified many other targets of miR-505-5p that may contribute to a nutrient sensing phenotype in the offspring of obese mothers. Among these are the SRCAP chromatin remodelling complex, which activates a transcriptional profile that shifts metabolism from anaerobic glycolysis to oxidative metabolism (61). Interestingly, the pSILAC revealed that many indirect targets of miR-505-5p (i.e. downregulated in pSILAC but do not contain miR-505-5p seed sequence) are related to FA sensing, and many of these also showed genetic associations with BMI in humans. Whilst we have focused on validating a few key direct target genes in the current study, it is unlikely that miR-505-regulates a complex process such as nutrient sensing via only one key target gene, and therefore it is likely that miR-505-5p controls the expression of several master regulators (e.g. transcription factors or chromatin remodelling complexes such as SRCAP) that modulate whole profiles of FA sensing genes. Notably, many of the proteins we found to be altered following over-expression of miR-505-5p are expressed in mitochondria, so future studies should aim to examine the effects of maternal obesity on offspring mitochondrial dynamics.

In conclusion, we have identified miR-505-5p as a novel regulator of FA metabolism pathways that is programmed in mice exposed to maternal obesity and could contribute to the increased HFD intake and adiposity seen in the offspring of obese mothers. This is consistent with previous animal studies showing a role for hypothalamic FA sensing in modulating food intake, and is corroborated by our human genetic data analysis. This study is one of the first reports defining a molecular route by which exposure to an altered nutritional environment *in utero* is able to alter feeding behaviour. Further defining the mechanisms underlying the programming of miR-505-5p by maternal obesity may hold promise for developing novel interventional strategies to prevent the adverse impacts of maternal obesity on offspring long-term health.

## 5. Authors contributions

LD and SEO conceived the project. ICF, LD, and SEO designed the experiments and interpreted the results. LD, ICF, LP, LWPW, DFT, JF, MC, and GB performed experiments. JD, AVP, LD and SEO supervised experiments. KAK, KKO and JRBP carried out the genetic analysis. LD wrote the manuscript. All authors reviewed and commented on the final version of the manuscript.

## Acknowledgments

We would like to thank Tom Ashmore, Claire Custance and Laura Hunter for assistance with animal husbandry, the Genomics and Transcriptomics Core for assistance with RNA-sequencing, and Dr. Robin Antrobus for assistance with mass spectrometry. Illustrative figures were generated using BioRender. This research was conducted using the UK Biobank Resource under application 9905.

## Supplementary Material Figure legends

Supplementary Table 1: A) Primer sequences of primers used in SYBR qPCR purchased from Sigma Aldrich B) Taqman Assay IDs of probes used in Taqman qPCR purchased from Thermofisher Scientific C) miRCURY LNA miRNA PCR assays purchased from Qiagen.

Supplementary Table 2: Significantly regulated miRNAs detected in paraventricular nucleus of the hypothalamus of offspring from obese mothers in miR-sequencing analysis.

Supplementary Table 3: Significantly regulated miRNAs detected in arcuate nucleus of the hypothalamus of offspring from obese mothers in miR-sequencing analysis.

Supplementary Table 4: Overlap of miR-505-5p targets with human genetic data on BMI variation and other functional datasets. Gene; gene symbol for human orthologue of mir505 target, chr; chromosome where Gene is locates, Summary; summarised count of supporting evidence across the four analyses, MAGMA P-value; p-value from the gene-level MAGMA test, MAGMA FDR-corrected P-value; MAGMA P-value after FDR correction, GWAS Signal; rsid for the GWAS signal proximal to Gene, position; chromosomal position of GWAS signal (GRCh 37), a1; effect allele, a0; alternate allele, freq1; observed frequency of a1, beta1; effect size estimate per copy of a1, se; standard error of beta1, p; p-value for the association, Is closest?; is Gene the closest gene to GWAS Signal, N ABC Enhancers; number of known ABC enhancers within the LD window (R-sq>0.8) of the GWAS Signal, P SMR; p-value for the SMR test, FDR-corrected P SMR; P SMR after FDR correction, P HEIDI; p-value for the Heidi test.

S1 Data: Original data for the graphs in Figure 1-5. Each tab includes data for the noted graphs in Figure 1-5. plus a tab for data mentioned in the text.

S2 Data: Raw data from p-SILAC experiment performed to identify changes in protein expression following miR-505-5p over-expression in GT1-7 cells.

