## supplementary data file for "Maternal obesity increases hypothalamic miR-505-5p expression in mouse offspring leading to altered fatty acid sensing and increased intake of high-fat food"

| Gene | Forward (5’-3’) | Reverse (5’-3’) |
| --- | --- | --- |
| Acot9 | gcgatacggctttggacctt | tccagactgtggatacgcct |
| Agpat3 | ctcacccctcttcagcttcg | gactccgaaggaagctgctc |
| Sdha | tcgacaggggaatggtttgg | taatcttccctggcatgggc |
| Slc25a10 | gcgggactacatgaccaagg | agagtagttgcgtcgttggc |
| Slc27a4 | cagccgggtcacaatgctac | tgaagacccggatgaaacgc |

A

B

| Gene | Assay ID |
| --- | --- |
| Atp11c | Mm01297974_m1 |
| Cpt1a | Mm01231183_m1 |
| Soat1 | Mm00486279_m1 |
| Sdha | Mm01352366_m1 |

C

| Gene | Assay ID |
| --- | --- |
| miR-505-5p | YP02104807 |
| Snord68 | YP00203911 |
| USP6 | YP02119464 |

**Supplementary Table 1:**

A) Primer sequences of primers used in SYBR qPCR purchased from Sigma Aldrich B) Taqman Assay IDs of probes used in Taqman qPCR purchased from Thermofisher Scientific C) miRCURY LNA miRNA PCR assays purchased from Qiagen.

|  | logFC | PValue |
| --- | --- | --- |
| mmu-miR-505-5p | 2.26790985 | 0.0101714 |
| mmu-let-7e-5p | 1.49047862 | 0.02151258 |
| mmu-let-7k | 1.418607 | 0.02855474 |
| mmu-miR-92b-5p | 1.01339219 | 0.04323239 |

**Supplementary Table 2:**

Significantly regulated miRNAs detected in paraventricular nucleus of the hypothalamus of offspring from obese mothers in miR-sequencing analysis

|  | logFC | PValue |
| --- | --- | --- |
| mmu-miR-505-5p | 3.15201921 | 7.38E-05 |
| mmu-miR-5105 | 1.30542539 | 0.00184717 |
| mmu-miR-6540-5p | 1.49630597 | 0.00273614 |
| mmu-let-7e-5p | 1.39679886 | 0.00330176 |
| mmu-miR-98-5p | 1.17038042 | 0.003708 |
| mmu-let-7f-5p | 1.26987087 | 0.00463843 |
| mmu-miR-6538 | 1.19286835 | 0.00473093 |
| mmu-let-7k | 1.31238993 | 0.00499318 |
| mmu-let-7c-5p | 1.29452908 | 0.00527431 |
| mmu-let-7d-5p | 1.37471051 | 0.00608837 |
| mmu-let-7a-5p | 1.21025834 | 0.008328 |
| mmu-miR-485-5p | 1.04037587 | 0.01619518 |
| mmu-miR-1224-3p | 1.10615633 | 0.02779869 |
| mmu-miR-383-5p | 1.1783129 | 0.03642917 |

**Supplementary Table 3:**

Significantly regulated miRNAs detected in arcuate nucleus of the hypothalamus of offspring from obese mothers in miR-sequencing analysis

| Gene | chr | Summary | BMI (UK Biobank + GIANT meta-analysis) | | | | | | | | | | | | | | |
| --- | --- | --- | --- | --- | --- | --- | --- | --- | --- | --- | --- | --- | --- | --- | --- | --- | --- |
|  |  |  | MAGMA P-value | MAGMA FDR-corrected P-value | GWAS Signal | position | a1 | a0 | freq1 | beta1 | se | p | Is closest? | N ABC Enhancers | P SMR | FDR-corrected P SMR | P HEIDI |
| CPT1A | 11 | 3 | 2.8E-05 | 2.7E-04 | rs497261 | 68192244 | T | C | 0.673 | 0.011 | 0.002 | 2.5E-08 | F | NA | 8.5E-06 | 2.1E-05 | 1.7E-03 |
| HSD17B12 | 11 | 3 | 3.2E-13 | 9.3E-12 | rs10838193 | 43898020 | G | A | 0.663 | 0.011 | 0.0018 | 1.7E-09 | F | 5 | 9.7E-26 | 4.9E-25 | 6.6E-04 |
| HINT1 | 5 | 3 | 9.7E-05 | 7.0E-04 | rs1363695 | 130378027 | C | T | 0.764 | 0.013 | 0.002 | 8.1E-10 | T | 1 | 8.4E-05 | 8.4E-05 | 2.0E-07 |
| SLC27A4 | 9 | 3 | 9.2E-07 | 1.3E-05 | rs7871866 | 131027982 | G | C | 0.840 | -0.018 | 0.0024 | 7.6E-14 | F | 19 | NA | NA | NA |
| SLC25A10 | 17 | 2 | 0.338 | 0.392 | rs11658335 | 80084821 | C | T | 0.418 | 0.011 | 0.0017 | 8.6E-11 | F | 17 | NA | NA | NA |
| EPHX1 | 1 | 2 | 3.6E-03 | 1.6E-02 | rs10915840 | 225668524 | G | A | 0.721 | 0.011 | 0.0019 | 1.9E-08 | F | NA | NA | NA | NA |
| HADHA | 2 | 2 | 0.666 | 0.689 | rs12468863 | 26940294 | C | T | 0.488 | 0.015 | 0.0016 | 6.6E-20 | F | 5 | NA | NA | NA |
| HSPA8 | 11 | 2 | 0.204 | 0.321 | rs7115089 | 122530591 | C | G | 0.628 | -0.011 | 0.0017 | 6.0E-11 | F | 4 | NA | NA | NA |
| AGPAT5 | 8 | 1 | 0.760 | 0.760 | NA | NA | NA | NA | NA | NA | NA | NA | NA | NA | 8.0E-05 | 8.4E-05 | 1.5E-02 |
| ERLIN1 | 10 | 1 | 0.055 | 0.146 | rs17094222 | 102395440 | T | C | 0.791 | -0.017 | 0.002 | 4.0E-18 | F | NA | NA | NA | NA |
| FDPS | 1 | 1 | 0.014 | 0.051 | NA | NA | NA | NA | NA | NA | NA | NA | NA | NA | 6.0E-05 | 8.4E-05 | 5.0E-02 |
| PITPNA | 17 | 1 | 0.276 | 0.348 | rs3923783 | 1843189 | C | A | 0.822 | 0.022 | 0.0022 | 4.1E-23 | F | NA | NA | NA | NA |
| ACAD9 | 3 | 1 | 0.172 | 0.295 | rs76594121 | 128189391 | T | G | 0.955 | 0.028 | 0.0047 | 1.6E-09 | F | NA | NA | NA | NA |
| ACADL | 2 | 1 | 0.019 | 0.062 | rs715 | 211543055 | T | C | 0.695 | -0.016 | 0.0019 | 1.3E-16 | F | NA | NA | NA | NA |
| AGPAT3 | 21 | 1 | 3.9E-03 | 1.6E-02 | NA | NA | NA | NA | NA | NA | NA | NA | NA | NA | NA | NA | NA |
| PAF1 | 19 | 1 | 1.8E-03 | 1.1E-02 | NA | NA | NA | NA | NA | NA | NA | NA | NA | NA | NA | NA | NA |
| DBI | 2 | 0 | 0.104 | 0.216 | NA | NA | NA | NA | NA | NA | NA | NA | NA | NA | NA | NA | NA |
| AGPAT4 | 6 | 0 | 0.524 | 0.563 | NA | NA | NA | NA | NA | NA | NA | NA | NA | NA | NA | NA | NA |
| HMGCL | 1 | 0 | 0.173 | 0.295 | NA | NA | NA | NA | NA | NA | NA | NA | NA | NA | NA | NA | NA |
| LPCAT3 | 12 | 0 | 0.267 | 0.348 | NA | NA | NA | NA | NA | NA | NA | NA | NA | NA | NA | NA | NA |
| CRAT | 9 | 0 | 0.076 | 0.178 | NA | NA | NA | NA | NA | NA | NA | NA | NA | NA | NA | NA | NA |
| ERLIN2 | 8 | 0 | 0.356 | 0.397 | NA | NA | NA | NA | NA | NA | NA | NA | NA | NA | NA | NA | NA |
| PTK2 | 8 | 0 | 0.080 | 0.178 | NA | NA | NA | NA | NA | NA | NA | NA | NA | NA | NA | NA | NA |
| SGPL1 | 10 | 0 | 0.222 | 0.321 | NA | NA | NA | NA | NA | NA | NA | NA | NA | NA | NA | NA | NA |
| SOD1 | 21 | 0 | 0.242 | 0.334 | NA | NA | NA | NA | NA | NA | NA | NA | NA | NA | NA | NA | NA |
| ACOT9 | X | 0 | NA | NA | NA | NA | NA | NA | NA | NA | NA | NA | NA | NA | NA | NA | NA |
| DHCR7 | 11 | 0 | 0.045 | 0.131 | NA | NA | NA | NA | NA | NA | NA | NA | NA | NA | NA | NA | NA |
| HADH | 4 | 0 | 0.338 | 0.392 | NA | NA | NA | NA | NA | NA | NA | NA | NA | NA | NA | NA | NA |
| PI4KA | 22 | 0 | 0.123 | 0.237 | NA | NA | NA | NA | NA | NA | NA | NA | NA | NA | NA | NA | NA |
| SOAT1 | 1 | 0 | 0.220 | 0.321 | NA | NA | NA | NA | NA | NA | NA | NA | NA | NA | NA | NA | NA |

**Supplementary Table 4:** Overlap of miR-505-5p targets with human genetic data on BMI variation and other functional datasets. Gene; gene symbol for human orthologue of mir505 target, chr; chromosome where Gene is locates, Summary; summarised count of supporting evidence across the four analyses, MAGMA P-value; p-value from the gene-level MAGMA test, MAGMA FDR-corrected P-value; MAGMA P-value after FDR correction, GWAS Signal; rsid for the GWAS signal proximal to Gene, position; chromosomal position of GWAS signal (GRCh 37), a1; effect allele, a0; alternate allele, freq1; observed frequency of a1, beta1; effect size estimate per copy of a1, se; standard error of beta1, p; p-value for the association, Is closest?; is Gene the closest gene to GWAS Signal, N ABC Enhancers; number of known ABC enhancers within the LD window (R-sq>0.8) of the GWAS Signal, P SMR; p-value for the SMR test, FDR-corrected P SMR; P SMR after FDR correction, P HEIDI; p-value for the HEIDI test.
